# Multi-omics analysis reveals discordant proteome and transcriptome responses in larval guts of *Frankliniella occidentalis* infected with an orthotospovirus

**DOI:** 10.1101/2023.10.31.564998

**Authors:** Jinlong Han, Dorith Rotenberg

**Affiliations:** North Carolina State University, Department of Entomology and Plant Pathology, Raleigh, North Carolina, 27695, U.S.A

**Author notes:** Correspondence: Dorith Rotenberg. Current address of J. Han: Department of Agricultural Biology, Colorado State University, Fort Collins, CO 80523, USA.

**Keywords:** western flower thrips, tomato spotted wilt virus, insect gut tissue, virus-vector interactions, weighted gene co-expression network analysis

## Abstract

The western flower thrips, *Frankliniella occidentalis*, is the principal thrips vector of *Orthotospovirus tomatomaculae* (order *Bunyavirales*, family *Tospoviridae*), a devastating plant-pathogenic virus commonly referred to as tomato spotted wilt virus (TSWV). The larval gut is the gateway for virus transmission by *F. occidentalis* adults to plants. In a previous report, gut expression at the transcriptome-level was subtle but significant in response to TSWV in L1s. Since it has been well documented that the relationship between the expression of mRNA and associated protein products in eukaryotic cells is often discordant, we performed identical, replicated experiments to identify and quantify virus-responsive larval gut proteins to expand our understanding of insect host response to TSWV. While we documented statistically-significant, positive correlations between abundance of proteins (4,189 identified) and their cognate mRNAs expressed in first and second instar guts, there was virtually no alignment of individual genes identified to be differentially modulated by virus infection at the transcriptome and proteome levels. Predicted protein-protein interaction networks associated with clusters of co-expressed proteins revealed wide variation in correlation strength between protein and cognate transcript abundance, which appeared to be associated with the type of cellular processes, cellular compartments, and network connectivity represented by the proteins. In total, our findings indicate distinct and dynamic regulatory mechanisms of transcript and protein abundance (expression, modifications, and/or turnover) in virus-infected gut tissues. This study provides molecular candidates for future functional analysis of thrips vector competence and underscores the necessity of examining complex virus-vector interactions at a systems level.

## 1. Introduction

Quantitative measurements of molecular species, such as RNA and protein, are essential for dissecting molecular mechanisms in disease biology. mRNA abundance in cells has been commonly used as a proxy for protein expression (Vogel and Marcotte, 2012), however, the accumulating body of evidence generated from animal systems with numerous genomics data sets clearly shows that the correlation of mRNA and protein levels varies significantly across cell types (Du et al., 2019) and transcript abundance insufficiently predicts protein abundance (Liu et al., 2016; Payne, 2015). This context-dependent and discordant association between mRNA and protein levels has been ascribed to a variety of complex regulatory mechanisms, including post-transcriptional and post-translational silencing, mRNA decay and protein degradation (Gygi et al., 1999; Vogel and Marcotte, 2012). As such, it is imperative to investigate both transcriptional and translational events and/or outcomes in order to gain more precise views of host cellular, tissue system and physiological responses associated with pathogen invasion.

*Orthotospovirus tomatomaculae*, in the genus *Orthotospovirus* (family *Tospoviridae*, order *Elliovirales*) and commonly known as tomato spotted wilt virus (TSWV), causes severe damage on many economically important food crops. It is transmitted by the western flower thrips, *Frankliniella occidentalis* (Pergande), in a circulative-propagative manner. The first (L1) and early second instar (L2) larval thrips are critical to the virus transmission cycle because adults - the most significant plant inoculators of TSWV during probing and feeding in the landscape - are only vector competent if they acquire TSWV as young larvae (Moritz et al., 2004; Van De Wetering et al., 1996). Upon entry of the L1 midgut, the virus replicates and spreads throughout surrounding visceral gut muscles and disseminates to the salivary glands during the late L2 stage. Virus is retained, albeit at lower abundance during the pupal stages (Schneweis et al., 2017), and when adults eclose, virus increases in the adult principal salivary glands (Montero-Astúa et al., 2016b). The virus is inoculated into the plant through thrips feeding (Montero-Astúa et al., 2016b; Ullman, 1992; Ullman et al., 1995, 1993). Virus infection and replication of this circulative virus in thrips offers opportunities for unraveling complex tissue response to infection. Considering the critical role of larval guts for successful establishment of infection, unveiling molecular responses to virus invasion of this tissue system is a significant step toward devising novel crop disease control strategies that disrupt the gut-virus interaction.

Numerous studies have utilized next-generation sequencing to investigate the molecular aspects of interactions between insect vectors and their associated plant viruses (Dietzgen et al., 2016; Ghosh and Ghanim, 2021; Heck, 2018; Rotenberg et al., 2015). However, only a handful of studies have examined the effect of plant virus infection on the gut of vector species (Brault et al., 2010; Geng et al., 2018; Han and Rotenberg, 2021; W. Zhao et al., 2016). Previously, we reported the first thrips gut transcriptome of *F. occidentalis* larvae in response to TSWV infection (Han and Rotenberg, 2021), and provided several candidate biomarkers of infection at a stage when TSWV is known to enter and invade the epithelium of the anterior midgut of L1 (*i.e.*, limited to a few cells) (Badillo-Vargas et al., 2019; Montero-Astúa et al., 2016b), and when virus titer is at its lowest during the transmission cycle (Badillo-Vargas et al., 2012; Han and Rotenberg, 2021; Rotenberg and Whitfield, 2018). It was the early L1 stage - three hours after a 24-hour exposure period on TSWV-infected plant tissue – that exhibited a relatively larger transcriptome-wide response to infection compared to the L2 stage (48 hours post exposure), when virus titers in the gut had increased 3-fold (Han and Rotenberg, 2021), and virus has disseminated and invaded other regions of the midgut (Montero-Astúa et al., 2016b).

This study attempts to gain a deeper understanding of the larval gut-TSWV interaction by i) quantifying modulation of the gut proteome by the virus; ii) identifying protein co-expression networks and functional enrichments represented by network members; and iii) examining the association between the gut proteome (this study) and transcriptome of TSWV-exposed and non-exposed larval thrips, as well as within network protein-transcript abundance correlations, by capitalizing on previously published gut transcriptome data. Collectively, our analyses revealed distinct responses of thrips gut tissues to virus infection at transcriptional and translational levels, stressing the importance of investigating virus-vector interactions using complementary approaches.

## 3. Results and Discussion

### 3.1. TSWV proteins detected in larval guts

Comparisons were made between the first instar larvae, three hours after a 24-hour exposure time to TSWV-infected plant tissue (3-L1-V) and second instar larvae ensuing from the 3-L1 cohort, 48 hours after removal from the infected plant tissue (48-L2-V). Structural and nonstructural proteins encoded by the five open reading frames of the TSWV genome were identified in the gut proteome. We documented a significantly greater normalized abundance of nucleocapsid protein (N, *P* = 0.005), a virion structural protein, and two non-structural viral proteins (NSs, *P* = 0.036 and NSm, *P* = 0.003), in 48-L2, compared to 3-L1 of virus-exposed insects (**Figure 1**). Virus accumulation in thrips guts after removal of the source of inoculum coupled with an increased abundance of non-structural proteins is indicative of virus replication (De Assis Filho et al., 2002; Ohnishi et al., 2001; Ullman et al., 1993). Increases in viral RNA (TSWV-N) from L1 to L2 bodies (Badillo-Vargas et al., 2012; Schneweis et al., 2017) or guts (Han and Rotenberg, 2021) of *F. occidentalis* during growth and development have been reported previously, corroborating our findings at the protein level.

**Figure 1.**
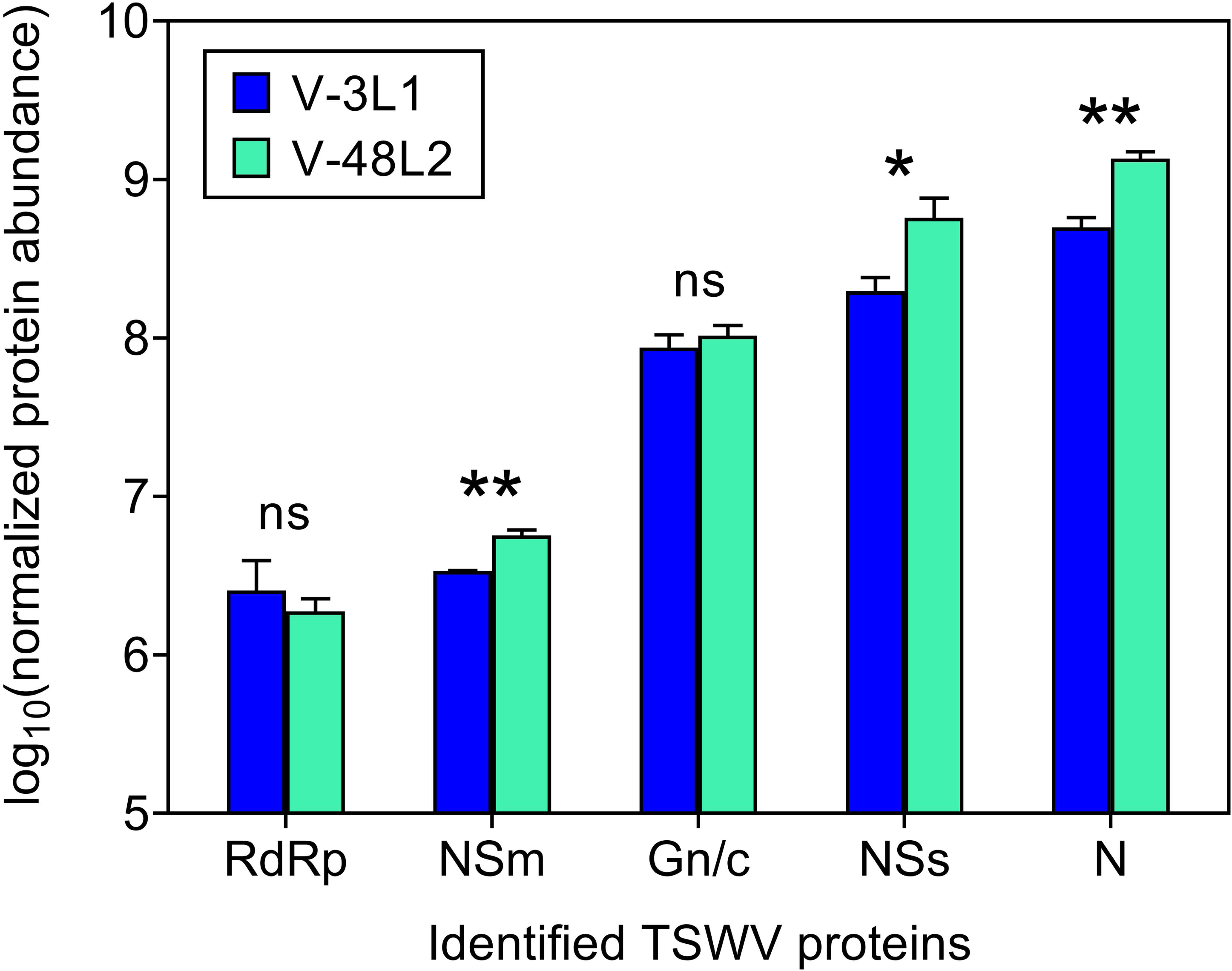
Accumulation of tomato spotted wilt virus (TSWV) proteins in *Frankliniella occidentalis* larval guts. One-way analysis of variance was performed for each protein to determine the effect of larval stage on log_2_-transformed, normalized abundance of TSWV proteins detected in the quantitative proteomics dataset: RNA-dependent RNA polymerase (RdRp), two non-structural proteins (NSm and NSs), glycoprotein precursor (GN_GC), and nucleocapsid protein (N). Error bar = standard error of the mean; ns = no significant difference. 3-L1 = first instar larvae, three hours after a 24-hour exposure time on TSWV-infected plant tissue; 48-L2 = second instar larvae ensuing from the 3-L1 cohort, 48 hours after removal from infected plant tissue.

While all attempts were made to avoid cross-contamination of virus-exposed (V) and non-exposed (NV) samples during tissue dissection and sample processing, there were peptides at low occurrence and low abundance that identified TSWV proteins in the NV-larval gut samples (**Supplementary File 1: Figure S1.A-B**). We examined the possible origin of these peptide sequences in the NV samples by i) performing NCBI blastp queries of the non-redundant protein database (Feb 13, 2024) using a low e-value cutoff (0.1) to rule out possible matches to other organismal proteins that may have been present in the gut samples, ii) performing local blastp queries to protein sequences retrieved from the NCBI Viral RefSeq Project (Bioproject PRJNA485481) for viromes identified in *F. occidentalis* (52 proteins) and *Thrips tabaci* (118 proteins) (Chiapello et al., 2021), including bunyaviruses, that may have been harbored by our thrips isolate, and iii) querying the *F. occidentalis* proteome with the identified TSWV peptides to rule out occurrence of identical thrips peptide sequences. Based on these *post priori* analyses, there was no apparent evidence of identical or similar peptide matches to other organisms or *F. occidentalis*. Of the 105 TSWV peptides identified in this study overall, 23 aligned to protein sequences of other bunyaviruses reported for thrips viromes. However, only ∼50% of TSWV peptide query lengths (7 – 32 amino acids), on average, aligned to any bunyavirus sequence match (N, NSs, NSm, or RdRP sequences). Moreover, 79% of the bunyavirus-aligned peptides were mismatched in annotation (e.g., N peptide with RdRP peptide). Since the NV sample peptides were identical in sequence and annotation to TSWV proteins, and only remotely aligned to other thrips-harbored bunyaviruses, we attribute their occurrence to some source or means of TSWV protein contamination during sample processing.

### 3.2. Influence of larval developmental stage and virus on normalized protein abundance in the gut proteome

Our proteomic study detected a total of 47,169 unique peptides across all gut samples, which identified 4,189 different *F. occidentalis* proteins. Principal component analysis (PCA) on a correlation matrix of normalized thrips protein abundance values by larval stage-treatment groups (NV and V) revealed that larval stages (3-L1 vs 48-L2), regardless of treatment, clustered separately along PC1, which explained the majority of the variation in the data set (35.37%), and that 3-L1 clustered by virus infection status (NV vs V) along PC2 (15.95% of the variation) (**Supplementary Figure S1A**). The finding that virus exposure and/or infection had a subtle effect on the gut proteome compared to the physiological status of the thrips vector is consistent with a report that *F. occidentalis* sex (male vs female adults) had a larger influence on the salivary gland proteome than TSWV infection of these adult glands (Rajarapu et al., 2022). Supporting the PCA observations, hierarchical clustering analysis revealed that samples grouped by both larval stage and virus infection status, further underscoring the dual influence of developmental stage and viral factors on the thrips gut proteome (**Supplementary Figure S1B**).

### 3.3. Differential abundance of larval gut proteins in response to virus

The larval gut response to TSWV significantly modulated the abundance of 233 and 121 proteins expressed in 3-L1 and 48-L2 guts, respectively (**Figure 2A**; *P* < 0.05 and at least 2-fold change compared to NV guts), referred herein as <u>v</u>irus-modulated, <u>d</u>ifferentially <u>a</u>bundant <u>p</u>roteins (vDAPs, **Supplementary File 1: Table S2**). Comparatively, the number of vDAPs in the *F. occidentalis* larval guts indicated a more robust gut response to TSWV exposure and infection in 3-L1 compared to 48-L2 (**Figure 2A**). This finding is consistent with our previous gut transcriptome study (Han and Rotenberg, 2021), where twice as many transcript sequences were differentially-abundant in guts of 3-L1 (105) compared to 48-L2 (49). Notably, virus exposure led to a greater apparent perturbation of gut proteins (relative numbers) compared to transcripts. The temporal proximity of exposure to and acquisition from the TSWV-infected plant host for the 3-L1 cohorts (3 hours post-removal of inoculum source) may, in part, explain the relatively larger response of L1s to the virus treatment. Within the time-frame of the present study, gut infection of young L1s is localized to epithelial cells of the anterior midgut, and within the L2 stage, TSWV has spread to other regions of the midgut and surrounding visceral muscles. This well documented phenomenon of tissue tropism (reviewed in Montero-Astúa, Stafford, et al., 2016), and the relative abundance of TSWV proteins in L1 compared to L2 guts (**Figure 1**) did not appear to explain the more robust effect of TSWV on L1s. We hypothesize that ingestion of infected plant tissue contributes to the larger gut response in L1s compared to L2s, which were sampled 48 hours after removal of the source of inoculum (∼24 hours post molting from L1 to L2). Examination of indirect (infected plant) vs. direct effects of TSWV on gut physiology is warranted to fully understand the nature of the virus-vector interaction.

**Figure 2.**
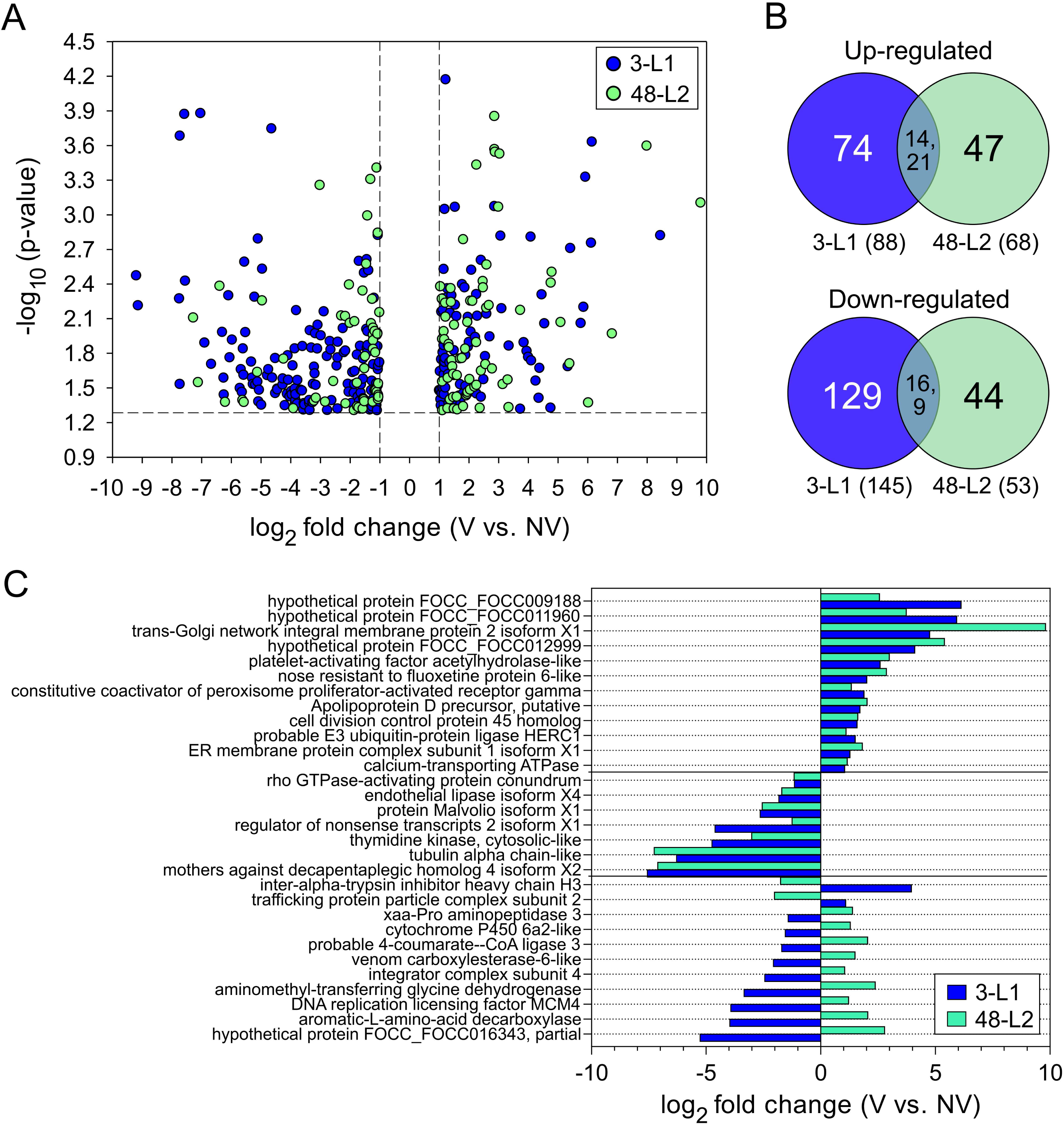
Differentially abundant proteins in *Frankliniella occidentalis* larval guts after exposure and infection by tomato spotted wilt virus (vDAPs). (A) Volcano plot showcasing the distribution of gut proteins by magnitude of change in normalized abundance of protein between virus (V) and no-virus (NV) conditions and level of significance of that change; proteins with a *p*-value < 0.05 and fold change ≥ 2 were identified as vDAPs. 3-L1 = first instar larvae, three hours after a 24-hour exposure time to plant tissue of the two conditions (V, NV); 48-L2 = second instar larvae ensuing from the 3-L1 cohort, 48 hours after removal from plant tissue. (B) Venn diagrams showing the numbers of up- and down-regulated vDAPs that are either shared or unique between the two larval stages. The values in parentheses indicate the total number of up- or down-regulated vDAPs in each larval stage. (C) Magnitude and direction of fold changes for vDAPs in common between larval stages.

A great number of vDAPs in 3-L1 guts (62%) were predominantly down-regulated (**Figure 2B**), as was reported for a whole-body proteome study of *F. occidentalis* L1s infected with TSWV (Badillo-Vargas et al., 2012). In contrast, the 48-L2 guts showed a relatively equal distribution of up- and down-regulated vDAPs. Of the 30 vDAPs determined to be shared between the two larval stages in the present study (**Figure 2B**, **Supplementary File 1:Table S3**), mothers against decapentaplegic homolog 4 (Smad4) and alpha-tubulin showed the most reduced abundances (**Figure 2C**) in both 3-L1 and 48-L2 guts. Smad4 is a signal transducer that mediates TGF-β signaling, regulating key genes involved in diverse physiological processes, including cell proliferation, homeostasis, differentiation, apoptosis, and immune response (Massagué, 2012). The significant down-regulation of Smad4 [log2 fold change (FC) ∼ -7] could disrupt TGF-β signaling pathways, which may have implications for gut immunity or stress responses during infection. TGF-β signaling is known to exert both suppressive and stimulatory effects on pathogen infections (reviewed in Deng et al., 2024). For example, studies have demonstrated that TGF-β1 enhances rubella virus and Zika virus infections in human cells through the Smad pathway (Pham et al., 2021; Trinh et al., 2022). However, the role of Smad4 in promoting or inhibiting TSWV infection in gut epithelial cells requires further investigation. Alpha-tubulin is a structural, cytoskeletal protein that forms microtubules (MT), adaptable tube-like structures that can rapidly reorganize, extend and depolymerize under different physiological states and external stimuli (Desai and Mitchison, 1997; Gasic et al., 2019). Depolymerization of microtubules (reviewed in Ohi et al., 2021) can lead to tubulin mRNA instability (Gasic et al., 2019) and subsequent reduced tubulin synthesis (Ben-Ze’ev et al., 1979), referred to as tubulin autoregulation. Autoregulation may have contributed to the reduced abundance of tubulins in guts of TSWV-infected L1s. Tubulin cytoskeletal networks in animal cells have been implicated in virus attachment to cell surfaces, intracellular trafficking, and virus replication (Naghavi and Walsh, 2017). More proximal to the present study, rice stripe virus (RSV), a circulative-propagated, plant-pathogenic RNA virus, was shown to upregulate and utilize alpha-tubulin for accumulation and dissemination along the route from gut cells to salivary glands of the small brown planthopper (Li et al., 2020). Negative regulation of tubulins in TSWV-infected L1 whole body proteomes (Badillo-Vargas et al., 2012) and the transcriptome of pro-pupae (P1) of *F. occidentalis* (Schneweis et al., 2017) is consistent with the present findings of larval gut tubulins. Other stage-shared proteins in the present study infer roles in DNA replication, transcription, intracellular transport, protein and mRNA turnovers, stress response, lipid, nucleotide, and amino acid metabolisms, detoxification, and cell cycle and structure (**Figure 2C, Supplementary File 1: Table S3**).

While many stage-shared vDAPs (63%) showed concordant direction of changes, others displayed opposite regulations between 3-L1 and 48-L2, which suggests a shift in the gut’s physiological or immune response as infection progresses (**Figure 2C**). For the most part, these shared proteins and their associated normalized abundances under the infection state indicate a negative regulation of gene expression and/or enhanced protein turnover in 3-L1 guts. However, this trend was inverted in 48-L2 guts, where most proteins were up-regulated. If and how TSWV modulates these cellular processes in thrips vector cells remains to be determined, and the current lack of a robust, immortalized thrips cell line for research beckons us to innovate ways to address these basic questions.

### 3.4. Gene ontologies (GO) and functional enrichment of larval gut proteins in response to TSWV

Of the 324 non-redundant vDAPs identified across the two larval stages, 79% were assigned GO terms (**Supplementary Figure S2, Supplementary File 1: Table S2**). The most predominant GO annotations in the biological process (BP) were ‘transport’ and ‘regulation of cellular process’ for 3-L1 and 48-L2 guts, respectively. In the molecular function, ’hydrolase activity’ was the most common annotation for both stages. The most frequent cellular compartments assigned to the vDAPs were ‘membrane’ for 3-L1 guts and ‘protein-containing complex’ for 48-L2 guts. Using STRING, it was determined that 82% and 73% of the thrips vDAP sequences for the 3-L1 and 48-L2 guts, respectively, identified orthologs (279 in total) in the annotated *D. melanogaster* proteome, and GO enrichment analysis of these orthologs revealed more granular, functional annotations encompassed by the mapped vDAPs (**Figure 3, Supplementary File 1: Table S4**). Not surprisingly, GO terms for the collection of vDAPs mirrored those of the larval stage overlap (DNA replication, nucleic acid metabolism, and cell structure) (**Figure 2C**).

**Figure 3.**
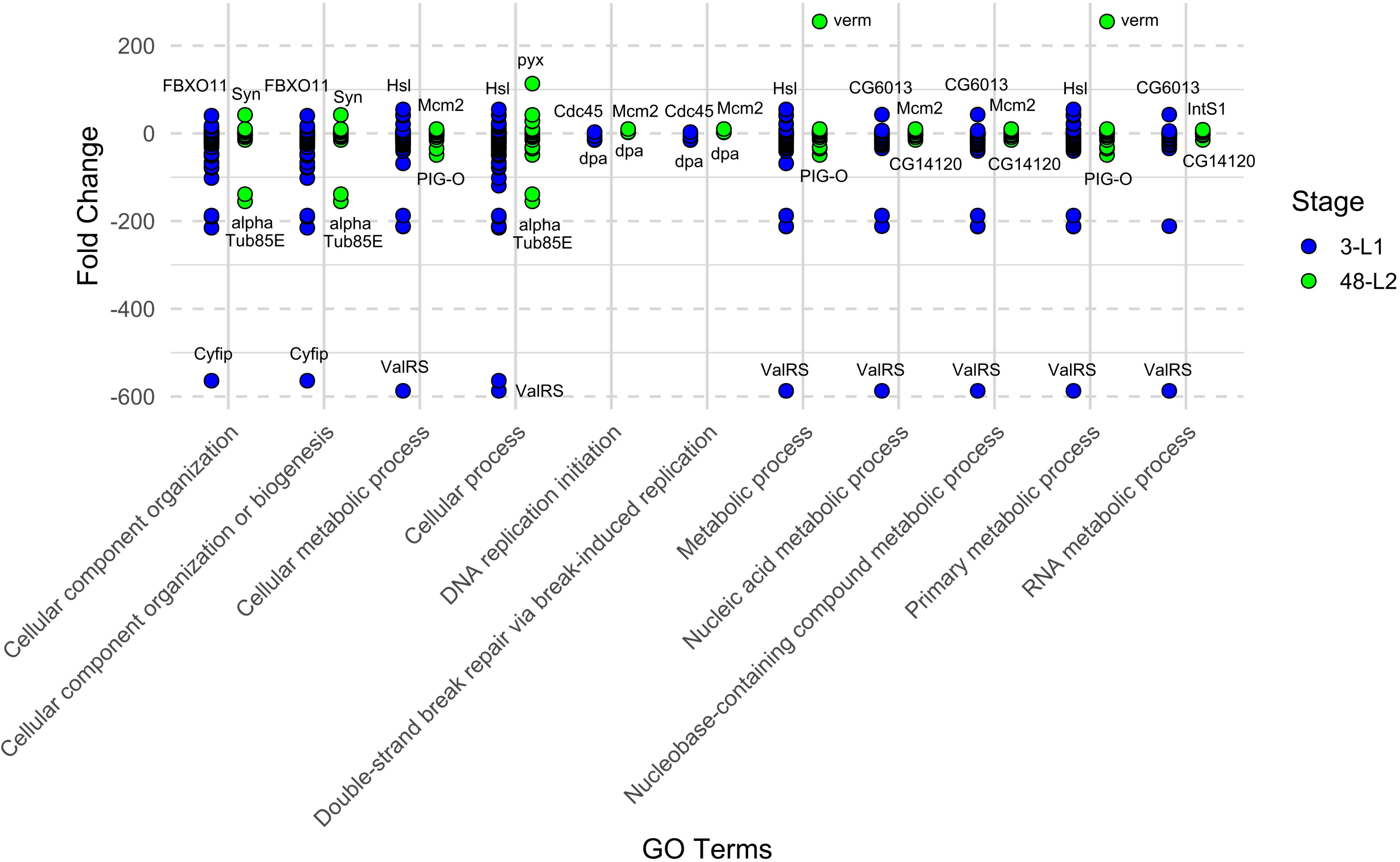
Fold change and gene ontology enrichment analysis of differentially abundant proteins in *Frankliniella occidentalis* larval guts after tomato spotted wilt virus exposure (V) and infection compared to the non-exposed (NV) condition (ie., vDAPs = 324 non-redundant proteins across the two larval stages); 3-L1 = first instar larvae, three hours after a 24-hour exposure time to plant tissue of the two conditions (V, NV); 48-L2 = second instar larvae ensuing from the 3-L1 cohort, 48 hours after removal from plant tissue. Seventy-nine percent of the vDAP dataset mapped to the *D. melanogaster* (Dm) proteome provided by the program. Enriched gene ontology (GO) terms in the biological process category are displayed, excluding obsolete terms using Revigo web tool; vDAPs with extreme fold change values within each stage and GO term are labeled with Dm ortholog notations: FBXO11 = F-box only protein 11; Cyfip = Cytoplasmic FMR1-interacting protein; Syn = Synapsin-1; alphaTub85E = Tubulin alpha-2 chain; Hsl = Hormone-sensitive lipase; ValRS = Valyl-tRNA synthetase, isoform C; Mcm2 = DNA replication licensing factor Mcm2; PIG-O = Phosphatidylinositol glycan anchor biosynthesis class O; pyx = Pyrexia; Cdc45 = Cell division cycle 45; dpa = Disc proliferation abnormal; verm = Vermiform; CG6013 = uncharacterized protein; CG14120 = uncharacterized protein; IntS1 = Integrator complex subunit 1.

Other enriched GO terms were revealed, namely ‘cellular component organization or biosynthesis’ and ‘primary metabolic process’ (**Figure 3**). We found that a vast majority of cellular component-associated vDAPs were down-regulated in larval gut tissues after virus infection. The 3-L1 guts exhibited a more pronounced cellular response than 48-L2 guts (65 vs 23 vDAPs). The most perturbed gut protein in this GO term was cytoplasmic FMR1-interacting protein-like isoform X1 (Cyfip, FC = -563.98), involved in actin cytoskeleton reorganization, mRNA translation, and endocytic trafficking and signaling (Abekhoukh et al., 2017; Napoli et al., 2008; Zhao et al., 2013). The substantial down-regulation of Cyfip in thrips guts may impair the cytoskeletal dynamics of gut epithelial cells, which are essential for maintaining gut structure and function. Among the up-regulated vDAPs, the F-box only protein 11 (FBXO11) showed a 40.41-fold increase in abundance in 3-L1 guts following virus infection. FBXO11 is a subunit of a SCF E3-ubiquitin ligase complex involved in degradation of diverse cellular proteins and has been implicated in silencing by si- and miRNAs at the step of RISC-loading in *Drosophila melanogaster* (Lee et al., 2009). More recently, it was reported that silencing of USB7 (a E3-ubiquitin ligase) in *F. occidentalis* adults resulted in reduced virus transmission to the host, but had no effect on acquisition (i.e., larval gut) (Shi et al., 2023).

In the enriched GO term of ‘primary metabolic process’, there were 83 and 40 vDAPs classified in the 3-L1 and 48-L2 guts, among which 66% and 48% were down-regulated, respectively (**Figure 3, Supplementary File 1: Table S4**). This category represented proteins with roles in protein ubiquitination or deubiquitination (proteolysis), DNA replication and repair, transcription regulation, regulation of mRNA splicing, ribosome biogenesis and translation, signal transduction, and lipid metabolism. vDAPs at the extremes in magnitude in change in response to TSWV for this GO term (**Figure 3**) were ValRS (FC = - 587.10), a valine-tRNA ligase known for its role in protein synthesis by incorporating valine into proteins, and verm (FC = 255.15), a chitin deacetylase involved in chitin remodeling and insect molting. Chitin is a key component of the cuticle and peritrophic membrane (PM) lining the midgut epithelium of many insects (Zhu et al., 2016). Cuticular proteins have shown to play an important role in virus transmission in diverse vector-borne plant virus systems. Unlike insects with PMs, hemipteran and thysanopteran midguts are lined with perimicrovillar membranes (PMMs) (Silva et al., 2004). Previously, Badillo-Vargas and Chen demonstrated that three cuticular proteins, containing chitin-binding domains, interacted directly with TSWV glycoprotein (G_N_) and were abundantly expressed in thrips guts and salivary glands. They hypothesized that chitin or chitin-like structures might be integrated into the PMM and salivary gland linings of thrips (Badillo-Vargas et al., 2019). Based on this hypothesis, we propose that the significant up-regulation of verm in 48-L2 guts may accelerate the conversion of chitin to a more soluble and less rigid chitosan polymer. This modification could potentially increase the permeability of the PMM lining, altering the gut environment in ways that may support TSWV replication or movement. In total, these vDAP identified orthologs with known, conserved functions with vital roles in cell biology and virology.

### 3.5. Larval gut co-expression protein networks

Using the collection of all *F. occidentalis* larval gut proteins identified in this study (4,189), WGCNA revealed a total of nine distinct clusters (i.e., modules) of co-expressed proteins grouped into four branches of the eigengene dendrogram, referred to as meta-modules (Langfelder and Horvath, 2008) (**Figure 4A**). vDAPs were distributed across eight modules, with the ‘turquoise’ module containing the most (112), 92% of which were down-regulated in response to TSWV (**Supplementary File 1: Table S2**). Modules that were positively correlated (**Figure 4B**, *P* < 0.05) with the TSWV condition were the ‘darkgreen (14% of vDAPs) and ‘darkgrey’ (11% of vDAPs) modules, and all of the vDAPs in these modules (87) were more abundant (up-regulated) in response to virus (3-L1 and 48-L2), with the exception of the sole two (down-regulated) associated with the 48-L2 stage (**Supplementary File 1: Table S2**). The ‘darkgreen’ and ‘darkgrey’ modules of proteins resided in two different meta-modules, suggesting different biological roles (Langfelder and Horvath, 2008) with regards to perturbation by the virus condition.

**Figure 4.**
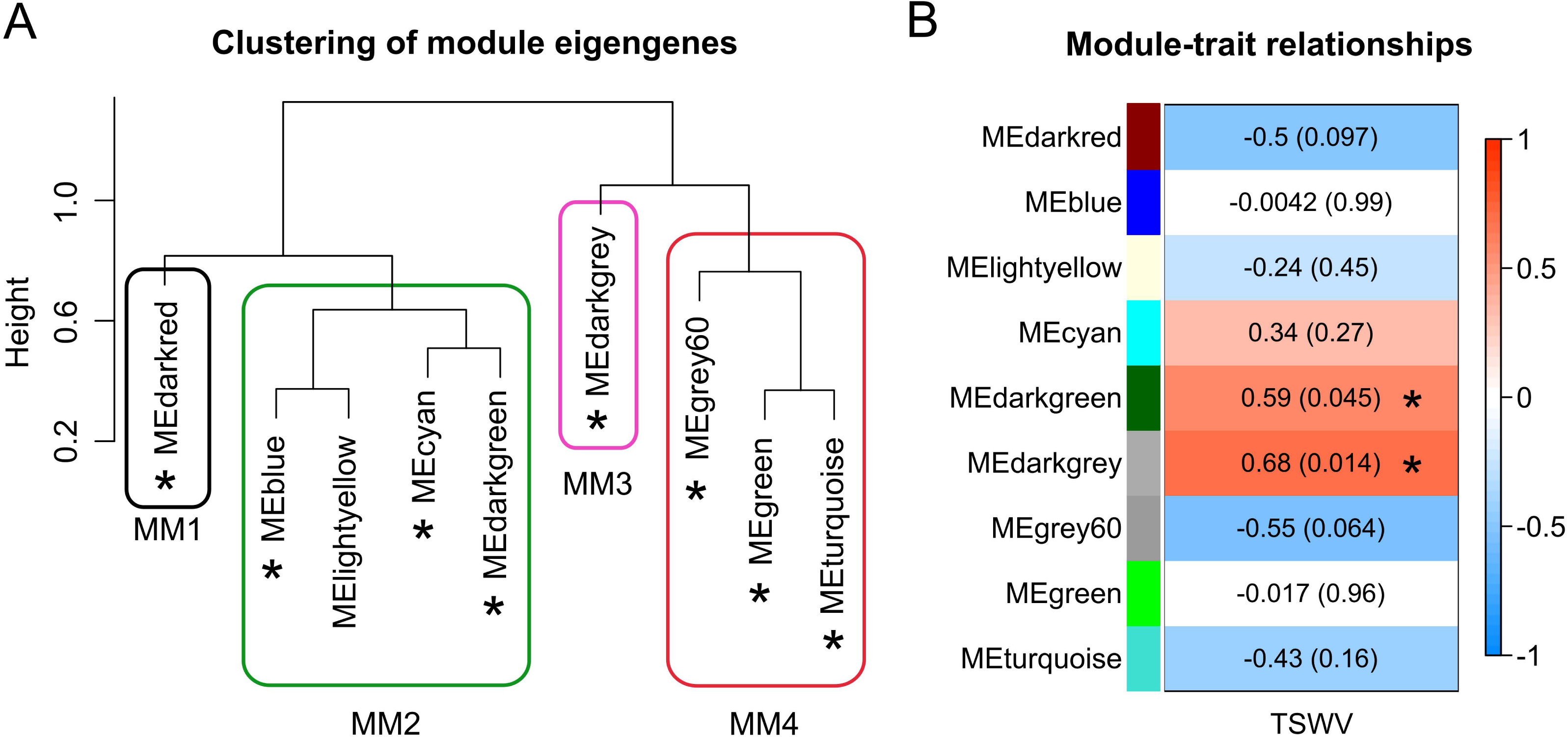
Weighted gene co-expression network analysis (WGCNA) identifies clusters of co-expressed larval gut proteins of *Frankliniella occidentalis.* A) The dendrogram illustrates intermodular relationships, with similar modules grouped together under the same meta-modules (MM); shown as branches on the eigengene dendrogram. Asterisks highlight modules that include sets of differentially abundant proteins in response to tomato spotted wilt virus exposure and infection (vDAPs). B) Module-trait relationships demonstrate significant correlations (*p-value* < 0.05) between modules (clusters of co-expressed proteins, represented as colored blocks on the left side of the column) and the external trait of tomato spotted wilt virus (TSWV) infection. WGCNA analysis was carried out using log_2_-transformed, normalized abundance values of all gut-expressed proteins, using samples from both virus-exposed (V) and non-exposed (NV) conditions with four biological replicates. Asterisks denote modules that exhibit a significant association with the external trait of TSWV infection.

Network membership analysis of the ‘darkgreen’ (192 proteins in total) and ‘darkgrey’ (420 proteins in total) modules identified 15 and 33 highly interconnected hub proteins, respectively (**Figure 5**), each with a module membership value (kME) no less than 0.86 and the greatest number of intramodular connections (kIN > 41) (**Supplementary File 1: Table S5**). All, except one, of the ‘darkgreen’ hub proteins were vDAPs (**Figure 5B)**, while only one of the ‘darkgrey’ hubs were responsive to TSWV under our experimental conditions (**Figure 5D**). Given that these hub proteins share the most connections with other co-expressed proteins (including other hubs), they likely play a critical role in regulating or coordinating activities of ‘neighborhood’ proteins in response to viral activities in larval gut cells.

**Figure 5.**
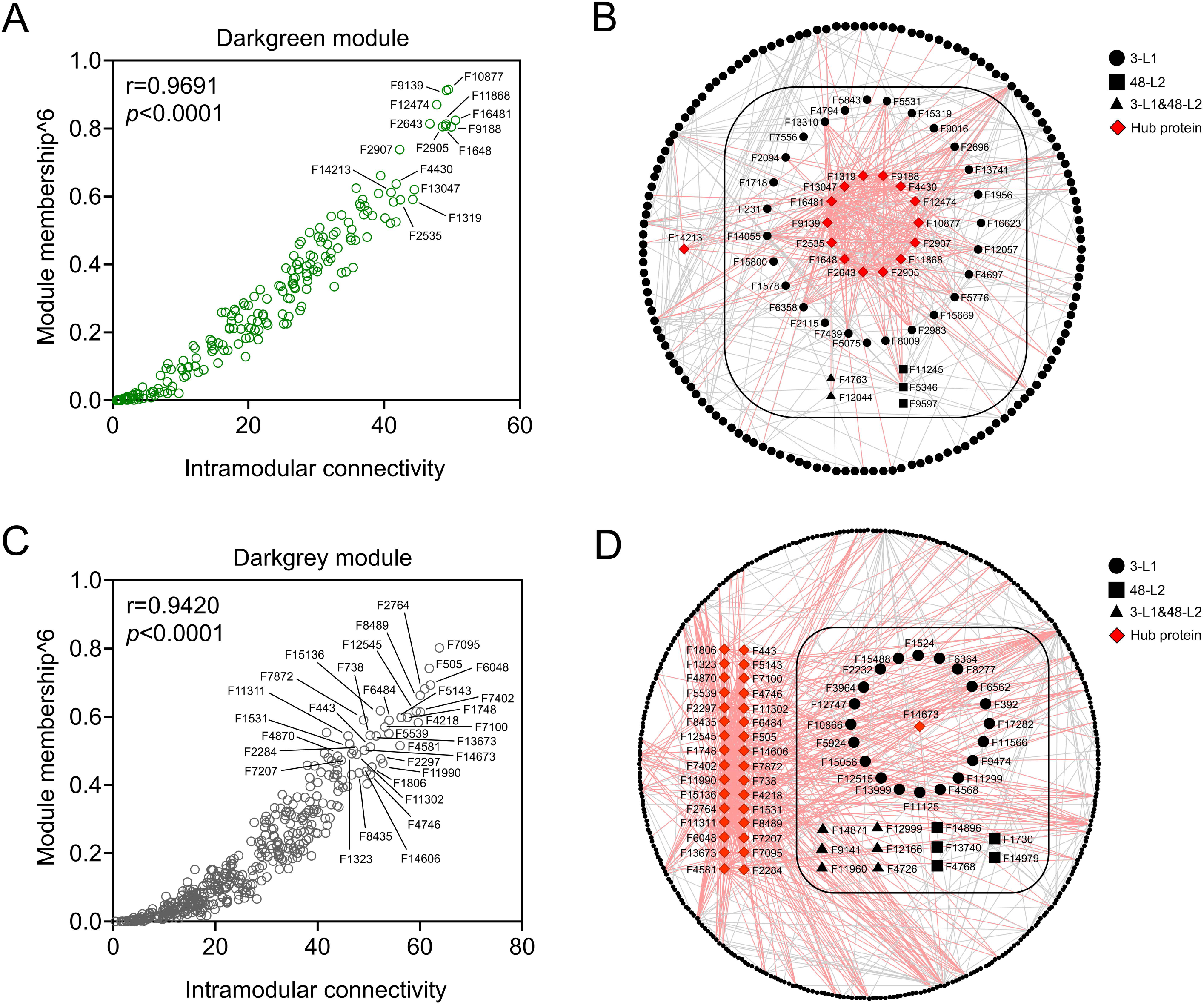
Intramodular network analysis of protein co-expression modules significantly correlated with the TSWV condition (see Figure 3B, darkgreen and darkgrey modules). Module membership (kME, correlation of a node to a module eigengene) and intramodular connectivity (kIN, sum of connection weights between a node and all its network neighbors) are plotted for all co-expressed proteins in the (A) darkgreen and (C) darkgrey modules. Pearson’s correlation coefficients (r) were calculated on the matrix of kME and kIN values for all proteins in each module. Proteins that ranked within the top 10% in both kME and kIN were considered candidate hub proteins (labeled data points). Intramodular relationships are visualized between co-expressed proteins in the (B) darkgreen and (D) darkgrey module networks. Proteins enclosed by the rounded square boxes represent tomato spotted wilt virus-modulated differentially abundant proteins (vDAPs) *Frankliniella occidentalis* larval guts. Hub proteins in each network are depicted as red rhombuses. Proteins shared between the 3-L1 and 48-L2 developmental stages, as well as those unique to each stage, are indicated by triangle, circle, and square shapes, respectively. For improved network visualization, a weight cutoff of ≥ 0.25 and ≥ 0.20 was applied in the darkgreen and darkgrey modules, respectively. Grey lines show connections between co-expressed proteins, while pink lines indicate hub-protein interconnections. 3-L1 = first instar larvae, three hours after a 24-hour exposure time to plant tissue; 48-L2 = second instar larvae ensuing from the 3-L1 cohort, 48 hours after removal from plant tissue.

Systems-level analyses of disease-associated omics datasets have led to the identification of key molecular pathways and potential new targets and biomarkers of diseases (Altaf-Ul-Amin et al., 2014). Functional analysis of the hub proteins and their connecting proteins with regards to thrips biology and thrips vector competence warrants further study. For example, in the present study, FOCC009188 (abbreviation: F9188), an uncharacterized protein with unknown function, was identified as a hub protein in the ‘darkgreen’ module. Notably, it was also shown as the most connected central hub gene in a co-expression network that was enriched with TSWV-responsive transcripts in *F. occidentalis* larval guts (Han and Rotenberg, 2021). Moreover, its protein abundance was significantly up-regulated in both 3-L1 and 48-L2 guts after virus infection, and its corresponding mRNA was previously shown to be ∼3-fold higher in TSWV-infected 3-L1 guts compared to uninfected controls.

### 3.6. Comparative analysis of the proteomes and transcriptomes of the larval gut of TSWV-exposed and non-exposed thrips

Identifiers (FOCC gene IDs) of all non-redundant larval gut proteins identified (4,189) were used to recover their associated transcripts and normalized abundance values from a previously published RNA-Seq expression data set for *F. occidentalis* larval transcriptomes under V and NV conditions (Han and Rotenberg, 2021) (**Supplementary File 1: Table S6**). To ensure the highest level of experimental consistency between the RNA-seq and proteomics analyses, our proteome experiment was meticulously designed and executed in strict accordance with the protocols defined in our RNA-Seq study. The search recovered transcript expression data for 99.3% of the protein data set. Statistical analysis of the paired transcript-protein expression data (log_2_ normalized abundance) revealed significant, positive correlations (Pearson, r) for guts of the 3-L1 (r = 0.625, *P* < 0.0001) and 48-L2 (r = 0.583, *P* < 0.0001) larval stages. Correlations of this strength and significance are remarkable for heterogeneous gut tissue (diverse cell types) that represent different regions of the insect gut (foregut, midgut and hindgut). Regression analyses on (x, y) pairs of TSWV-modulated proteins for each virus condition (NV/V) revealed a more significant, yet weak association between protein and transcript abundance in V-3-L1 guts (R^2^ = 18.1, *P* < 0.0001, n = 228) compared to NV-3-L1 (R^2^ = 0.11, *P* = 0.265, n = 228) guts (**Figure 6B vs 6A**). Despite significant positive correlations in larval guts in general, use of transcript abundance to explain the variation in protein abundance had limited value for TSWV-modulated proteins.

**Figure 6.**
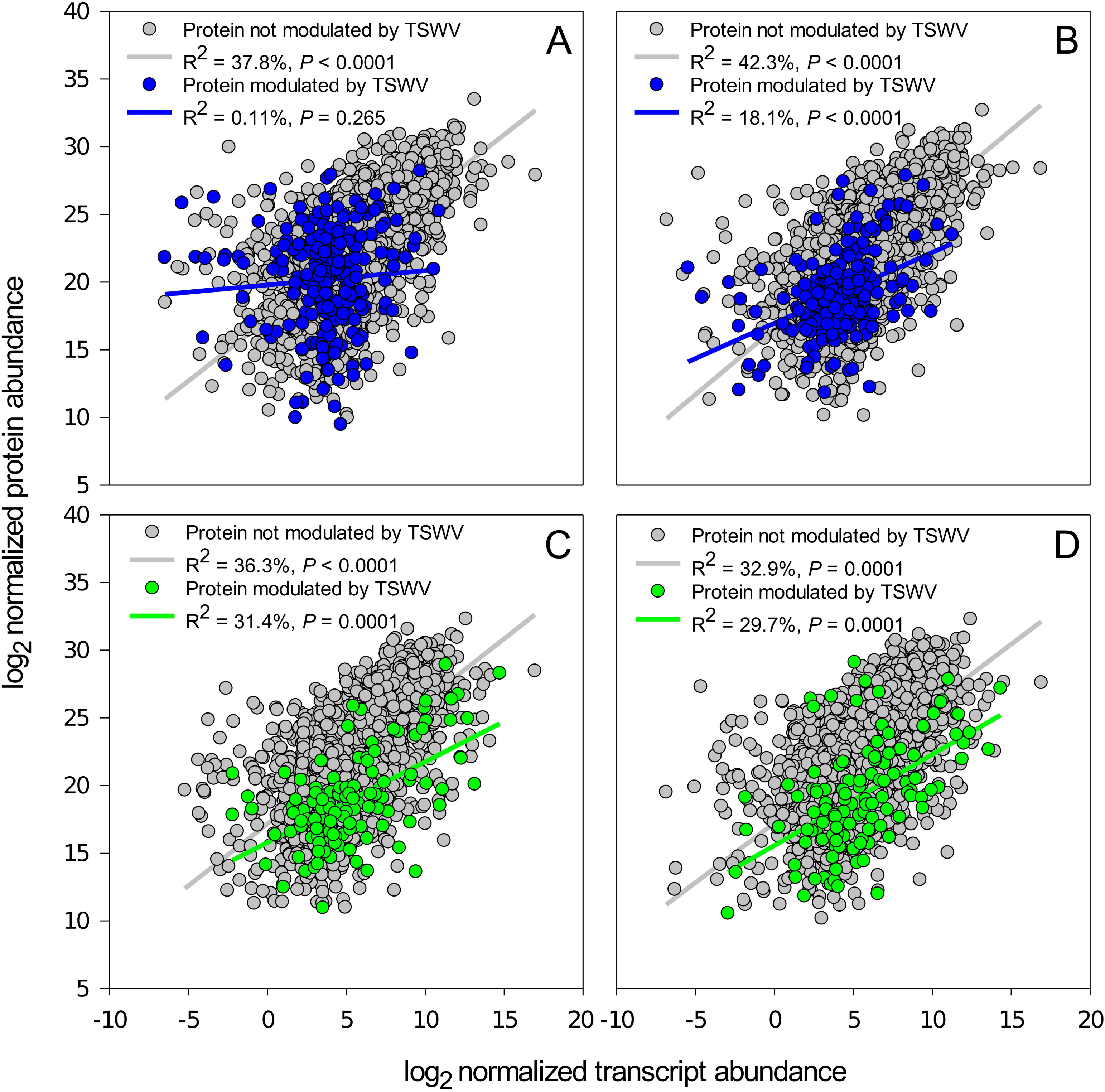
Correlations between the abundance of *Frankliniella occidentalis* larval gut proteins and their cognate transcripts for (A) first instar larvae, 3 hours after a 24-hour exposure time on healthy plant tissue (NV 3-L1); (B) first instar larvae, 3 hours after a 24-hour exposure time on TSWV-infected plant tissue (V 3-L1); (C) second instar larvae ensuing from the NV 3-L1 cohort, 48 hours after removal of plant tissue (NV 48-L2); and (D) second instar larvae ensuing from the V 3-L1 cohort, 48 hours after removal of infected plant tissue (V 48-L2). Pearson’s correlation and linear regression analyses were performed on log_2_-transformed normalized abundances values for proteins and their cognate transcript. Blue- and green-colored data points indicate TSWV-modulated, differentially-abundant proteins (vDAPs). Gray and blue/green lines depict best-fit regression lines for the indicated subsets of proteins.

The two ‘omics expression datasets were cross-queried (by OGS IDs) to identify overlapping sequences that were significantly (*P* < 0.05) perturbed by TSWV as compared to NV counterparts. In each stage, only two sequences were shared between the data sets, with all agreeing in the direction of changes in both protein and transcript levels (**Table 1**). Among them, FOCC000766 was further shared between 3-L1 and 48-L2 guts and consistently down-regulated in both stages. This sequence was annotated as phospholipase A1-like (PLA1), which functions in lipid metabolism, signal transduction, and membrane remodeling (Richmond and Smith, 2011). Down-regulation of PLA1 activity may impact lipid remodeling and cellular membrane properties, facilitating virus infection and dissemination in gut epithelium. The other two sequences were uncharacterized proteins with unknown functions (**Table 1**); however, FOCC009188 stands out as a particularly strong candidate for further investigation due to its consistent responses to virus infection and intramodular hub centrality within co-expressed protein and transcript modules (Han and Rotenberg, 2021) (**Figure 5B**).

**Table 1.** Shared TSWV-responsive larval gut proteins and transcripts identified from the proteomics and transcriptomics analyses.

| Stage<sup>x</sup> | ID | vDAPs<sup>#</sup> |  |  | DETs<sup>&</sup> |  |  | Gene description |
| --- | --- | --- | --- | --- | --- | --- | --- | --- |
| | | FC<sup>\$</sup> | P-value | Regulation | FC | P-value | Regulation | |
| 3-L1 | FOCC009188 | 69.73 | 0.002 | Up | 3.18 | 0.008 | Up | hypothetical protein FOCC_FOCC009188 |
|  | FOCC000766 | -3.57 | 0.023 | Down | -4.23 | 0.000 | Down | phospholipase A1-like |
| 48-L2 | FOCC001083 | -18.89 | 0.018 | Down | -8.75 | 0.006 | Down | hypothetical protein FOCC_FOCC001083 |
|  | FOCC000766 | -3.27 | 0.047 | Down | -5.86 | 0.021 | Down | phospholipase A1-like |

<sup>x</sup> developmental stage: 3-L1 = first instar larvae, 3 hours after a 24-hour exposure time on non-infected or TSWV-infected plant tissue; 48-L2 = second instar larvae ensuing from the 3-L1 cohorts, 48 hours after removal of non-infected or infected plant tissue; <sup>#</sup> vDAPs: virus-responsive, differentially abundant proteins identified from this study; <sup>&</sup> DETs: differentially-expressed transcripts from a previous gut transcriptome study (Han and Rotenberg, 2021); <sup>\$</sup> FC: fold change in normalized abundance between virus (V) and no-virus (NV) treatments.

Among the 324 non-redundant vDAPs, 99% displayed unaltered mRNA abundances after virus exposure, implying undisturbed transcription while post-transcriptional processes influenced translation efficiency (Sunnerhagen, 2007; B. S. Zhao et al., 2016). This gave rise to variable translation, with some proteins being enhanced and others diminished. Similarly, among the 147 non-redundant, differentially-expressed transcripts from our previous gut transcriptome study (Han and Rotenberg, 2021), 98% maintained stable protein levels despite potential fluctuations in mRNA levels. This protein stability may arise from adjustments in translation rates, changes in mRNA stability, and post-translational modifications like phosphorylation, ubiquitination, or acetylation, ensuring protein constancy even in the face of shifting mRNA abundances (Deribe et al., 2010; Mata et al., 2005; McCarthy, 1998; Pradet-Balade et al., 2001). Collectively our data indicates distinct transcriptional and translational responses of the larval gut to TSWV infection, given that consistent experimental conditions, parameters, and timing were followed intentionally for the two ‘omics studies. These distinct responses likely reflect temporally-dynamic and direct physical and biochemical interactions between the bunyavirus and host cell machinery, molecular building blocks (nucleotides and amino acids), and myriad unknown host factors available in the cytoplasm to complete its ‘life-cycle’ [transcription, translation, replication, virion assembly] (reviewed in Elliott & Schmaljohn, 2013). TSWV-induced, post-translational modifications of host proteins (Gong et al., 2023) may also contribute to the discordance between transcriptomic and proteomic response. Mass spectrometry-based, discovery proteomics methodologies, such as the one used in the present study, are not designed to handle identification of PTMs (Kim et al., 2016), especially with the use of a reference proteome database composed of single protein models per gene model. Therefore, use of existing RNA-Seq datasets for *F. occidentalis* in public repositories to identify transcript variants and isoforms for gene models and incorporation of PTM detection techniques for new proteome datasets may provide a more complete view of host response to virus or other persistent perturbations.

### 3.7. Intramodular analysis of correlations between protein and transcript abundance, and functional enrichment

In an attempt to approach the basis of correlations between transcriptome and proteome expression patterns in larval guts, rank correlation coefficients (Spearman’s rank coefficient, ⍴) were calculated for protein-transcript pairs for the nine sets of module members for each larval stage-treatment combination. The analyses revealed significant, positive associations (⍴) for all the nine modules. However, the strength of the association varied considerably among these modules (**Figure 7**).

**Figure 7.**
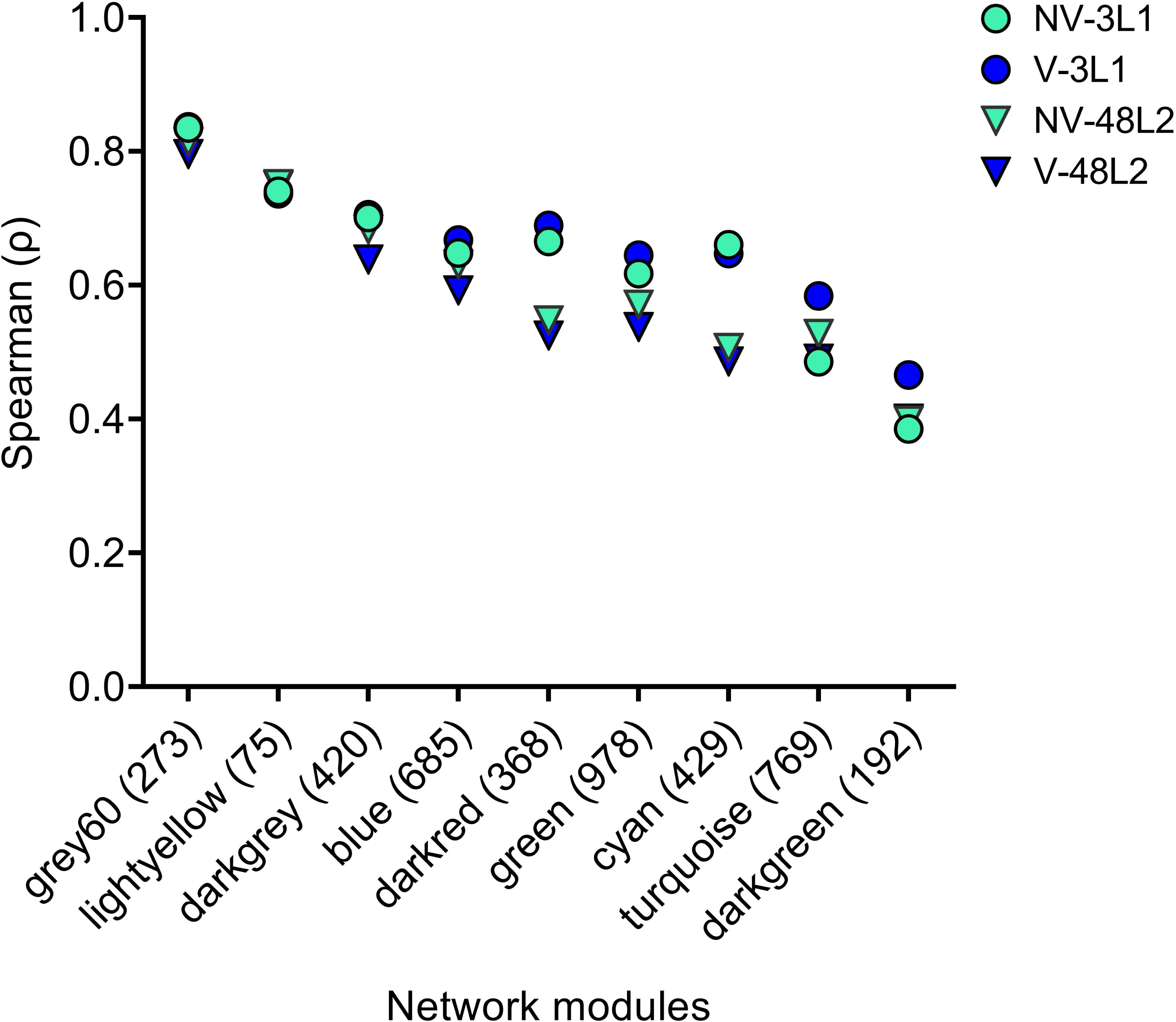
Associations between larval gut proteins and their cognate transcripts in *Frankliniella occidentalis* for the nine network-modules of co-expressed proteins. Spearman’s rank correlation coefficients (ρ) were calculated on log_2_-transformed, normalized abundance of proteins and transcripts each larval stage - virus condition combination for each of the network modules. The number within parentheses () indicates the number of proteins within each module. V/NV = virus-exposed (and infected)/non-exposed; 3-L1 = first instar larvae, three hours after a 24-hour exposure to the condition (V or NV); 48-L2 = second instar larvae ensuing from the 3-L1 cohort, 48 hours post exposure to the condition.

Functional enrichment analyses of protein-protein interactions (PPI) in five module networks were performed, one of which represented the highest strength in protein-transcript associations in the data set [grey60 (mean ⍴ = 0.821)], the two [darkgrey (mean ⍴ = 0.682) and darkgreen (mean ⍴ = 0.413)] that were significantly (positive) associated with TSWV (**Figure 4B**), and the two [darkred (mean ⍴ = 0.607) and cyan (mean ⍴ = 0.575)] that showed distinct protein-transcript correlation strengths between 3-L1 and 48-L2 guts (**Figure 7**). A summary of the functional enrichments and associated metrics for these five networks is presented in **Supplementary File: Table S7.** The <u>grey60</u> network (232 nodes) was enriched (*P* < 1x10^-16^) in ribosomal proteins and other highly conserved translational machinery and translation factors (e.g. eukaryotic elongation factors). The <u>darkgrey</u> network (346 nodes) was enriched (*P* < 1x10^-16^) in proteins associated with mitochondrial structure and processes, including pyruvate metabolism and ATP synthesis (pyruvate dehydrogenase complex), electron transport and energy production (iron-sulfur cluster binding), and fatty acid beta-oxidation and protein phosphorylation. The <u>darkgreen</u> network (145 nodes) had relatively fewer member connections (no. of edges in relation to no. of hubs) compared to the other modules of comparable neighborhood size, and functional enrichment was statistically supported, albeit at a lower significance (*P* < 1.8^-3^). It was enriched for functions associated with the TRAPPIII protein complex, a key regulator of intracellular membrane trafficking from the endoplasmic reticulum to the Golgi apparatus, particularly in the secretory pathway and autophagy. The <u>darkred</u> network (305 nodes) was enriched (*P* < 1x10^-16^) in proteins related to mitochondrial processes, fatty acid metabolism, proteasome activity, and virus entry regulation. The <u>cyan</u> network (349 nodes) was enriched (*P* < 1x10^-16^) in proteins involved in processes, such as mitochondrial fission, fatty acid elongation, proteasome degradation, and storage proteins. The darkred network features a more prominent role for clathrin and viral entry, suggesting a potential involvement in trafficking and virus-host interactions, whereas the cyan network emphasizes larval serum protein complexes and ER-associated processes, indicating a possible role in protein storage and degradation. Both networks share a common theme: protein degradation, specifically through the proteasome. This suggests that protein turnover is a critical process in both 3-L1 and 48-L2 guts. However, the specific proteins targeted for degradation may differ between the two stages as evident by the reduced correlations between mRNA and protein levels in 48-L2 guts (darkred mean ⍴ = 0.537, cyan mean ⍴ = 0.497) compared to 3-L1 guts (darkred mean ⍴ = 0.678, cyan mean ⍴ = 0.654) (**Figure 7**).

WGCNA-derived modules represent different sets of interconnected genes, and by virtue of these connections, indicate coordinated interactions that culminate into distinguishable cellular processes. Based on the intramodular correlations between protein and transcript abundance and functional enrichments determined by our analyses, we posit that the strength of the transcript-protein association may be explained, in part, by the type of cellular processes and cellular compartmentalization represented by the module (Du et al., 2019; Tasaki et al., 2022), and possibly connectivity (number of connections) of a protein with other proteins within the PPI network, like conserved translational and post-translational machinery (e.g., ribosomal protein complexes, proteasome) (Tasaki et al., 2022). Indeed, the enrichment of the grey60 and darkgrey networks with translation elongation factor complexes and mitochondrial compartmentalized proteins, respectively, is consistent with this hypothesis, as is the darkgreen network with fewer PPI edges and enrichment in proteins that play roles in intracellular membrane trafficking and autophagy. One study addressed the discordance of transcript to protein abundance by training models on cancer tumor ‘omics datasets and validating the predictive power of non-cognate transcript abundance on proteins in large and small multiprotein complexes associated with diverse biological processes (Srivastava et al., 2022). Determination of transcript variants and post-translational modifications in tissues of *F. occidentalis* and/or other thrips vectors of orthotospoviruses may provide a pathway towards identification of reliable biomarkers of vector competence, significantly advancing our understanding of virus transmission by thrips vectors.

## 3. Experimental Procedures

### 3.1 Biological source for larval gut protein samples

The source of *F. occidentalis* (lab colony on green bean pods), isolate of TSWV (*Emilia sonchifolia* lab culture), and the TSWV acquisition protocol – i.e., age-synchronized L1 cohorts subjected to a 24-hour acquisition access period (AAP) on infected (V) or non-infected (NV) plant tissue, then moved to green beans for gut clearing and development - were prepared and used precisely as described for our larval gut transcriptome study (Han and Rotenberg, 2021). The gut proteome experiment consisted of the two treatments (V and NV) and two sampling time-points post AAP [3-hrs (L1 stage) and 48-hrs (L2 stage), and the experiment was conducted three independent times (i.e., biological replicates) (**Supplementary Figure S3**). Infection rates of larval cohorts (V and NV) were determined by arbitrarily subsampling 10 individuals (L1) and subjecting each larval body to RNA extraction and cDNA synthesis followed by real-time PCR detection of TSWV nucleocapsid (N) RNA and *F. occidentalis* actin RNA (internal reference) as described previously (Han and Rotenberg, 2021). Based on this strategy, it was confirmed that each V cohort (all three bio-replicates) exhibited a 100% infection rate, and TSWV N RNA was not detected in the NV cohorts.

### 3.2. Thrips gut dissection

To ensure the highest level of experimental consistency for comparing proteome expression data to the published RNA-seq dataset (Han & Rotenberg, 2021), we executed experiments and thrips dissections in strict accordance with the protocols that formed the basis for the RNA-seq study, including taking into account possible effects of diurnal cycles on gene expression (e.g., all gut dissections performed at same period during the day). Larval dissections were performed to carefully remove gut tissue, which included the foregut, midgut, and partial hindgut as described previously (Han and Rotenberg, 2021), with adaptations for downstream protein analyses. A detailed experimental design is illustrated in **Supplementary Figure S3** in the current study. Each thrips larva was placed on a sterile microscope slide, immersed in ice-cold phosphate-buffered saline (PBS) buffer (∼80 µl), then decapitated using a sterile, polymer-coated scalpel blade (#15, Southmedic Inc., Barrie, Ontario, Canada). Undesired tissues, including principal and tubular salivary glands, fat body, and other residual body parts were carefully removed from the gut tissue using an ultrafine single deer hair tool (Ted Pella, Redding, CA). After dissection, an additional ice-cold PBS buffer (∼100 µl) was added to the slide to further wash away the unwanted residual tissue and body contents from the gut tissue. Each gut was immediately pulled through the surface of the buffer with the single deer hair tool and transferred into 200 µl of fresh ice-cold PBS buffer in a sterile 1.5-ml microfuge tube. This process was repeated to pool a total of 100 guts in the same tube, which was placed on ice throughout the entire process. Each replicate of the experiment was performed under the same laboratory conditions, during the same time period of the day, and by alternating between 20 NV-larvae and 20 V-larvae until obtaining 100 guts for both samples (using separate tools for each treatment). Sample tubes containing gut tissues were centrifuged at 8,000 g for 5 minutes and excess PBS buffer was carefully removed to achieve 20 µl in each tube. Gut samples were immediately flash-frozen in liquid nitrogen and stored at -80℃ until use. A total of 12 gut samples were prepared (two treatments, two sampling time points, three bio-replicates). To ensure that 100 guts would provide sufficient protein concentrations for downstream processing, we performed the bicinchoninic acid assay (BCA) (Thermo Fisher Scientific, Waltham, MA) to estimate total protein concentration on an entirely separate pool of 100 L1 guts.

### 3.3. Tissue-processing and ultra-performance liquid chromatography-tandem mass spectrometry analysis

The 12 frozen gut samples were submitted to the Duke Proteomics and Metabolomics Core Facility for sample processing and ultra-performance liquid chromatography-tandem mass spectrometry (UPLC-MS/MS) analysis and mass spectrometry data acquisition using state-of-the-art proteomic tools, resources and expertise. Briefly, each sample was spiked with undigested bovine casein as an internal quality control standard, which were then supplemented with 100 µl of 8M urea/50 mM ammonium bicarbonate. Proteins were solubilized by probe sonication (30% power, 3X). Protein content was estimated based on the protein concentration determined by BCA assay, and 4.5 ng of protein extract was used for enzymatic digestion with 20 ng/µl sequencing grade trypsin (Promega, Madison, WI) for 1 hour at 47°C, and eluted using 50 mM TEAB, followed by 0.2% FA, and lastly using 50% ACN/0.2% FA. All samples were then lyophilized to dryness and resuspended in 12 µl of 1% trifluoroacetic acid/2% acetonitrile with 12.5 fmol/µl of yeast alcohol dehydrogenase (ADH). A study pool quality control (SPQC) was created by combining equal volumes of each sample.

Quantitative LC/MS/MS was performed with 4 µl of each sample using a nanoAcquity UPLC system (Waters Corp) coupled to a Thermo Orbitrap Fusion Lumos high resolution accurate mass tandem mass spectrometer (Thermo) equipped with a FAIMSPro device via a nanoelectrospray ionization source. Samples were randomized across all four unique biological groups with sample pool QC standards run periodically throughout the study window. This acquisition strategy is, in part, designed to access a “real signal” across a large sample cohort, and provides support that the signals are not an artifact of the analytical system or experimental design. Briefly, the sample was first trapped on a Symmetry C18 20 mm×180 µm trapping column (5 µl/min at 99.9/0.1 v/v water/acetonitrile), after which the analytical separation was performed using a 1.8 µm Acquity HSS T3 C18 75 µm×250 mm column (Waters Corp.) with a 90-min linear gradient of 5 to 30% acetonitrile with 0.1% formic acid at a flow rate of 400 nl/min with a column temperature of 55℃. Data collection on the Fusion Lumos mass spectrometer was performed for three different compensation voltages (CV) (-40v, -60v, -80v). Within each CV, a data-dependent acquisition (DDA) mode of acquisition with a r=120,000 (@ m/z 200) full MS scan from m/z 375-1500 with a target AGC value of 4e5 ions was performed. MS/MS scans were acquired in the linear ion trap in “rapid” mode with a target AGC value of 1e4 and max fill time of 35 milliseconds. The total cycle time for each CV was 0.66 seconds, with total cycle times of 2 seconds between full MS scans. A 20-second dynamic exclusion was employed to increase depth of coverage. The total analysis cycle time for each sample injection was approximately 2 hours.

### 3.4. Mass spectrometry data acquisition and analysis

Following 16 total UPLC-MS/MS analyses (including four replicative SPQC samples), data were imported into Proteome Discoverer 2.4 (Thermo Scientific Inc.) and individual UPLC-MS/MS data files were aligned based on the accurate mass and retention time of detected precursor ions (“features”) using Minora Feature Detector algorithm in Proteome Discoverer. Relative peptide abundance was measured based on peak intensities of selected ion chromatograms of the aligned features across all runs. The MS/MS data was searched against the combined protein sequence database consisting of the *Frankliniella occidentalis* protein sequence database (Rotenberg et al., 2019), tomato spotted wilt virus protein sequences [AAF80981.1 (G_N_/G_C_ glycoprotein precursor); AAL55403.1 (RNA-dependent RNA polymerase); in house clones for NSs, NSm, and N]; a common contaminant/spiked protein database (bovine albumin, bovine casein, yeast ADH, etc.), and an equal number of reversed-sequence “decoys” for false discovery rate (FDR) determination. Mascot Distiller and Mascot Server (v 2.5, Matrix Sciences) were utilized to produce fragment ion spectra and to perform database searches. Database search parameters included fixed modification on Cys (carbamidomethyl) and variable modification on Met (oxidation). Search tolerances were 2 ppm precursor and 0.8 Da product ion with full trypsin enzyme rules. Peptide Validator and Protein FDR Validator nodes in Proteome Discoverer were used to annotate the data at a maximum 1% protein FDR based on q-value calculations. A peptide matched to multiple different proteins was exclusively assigned to the protein that had more identified peptides. Protein homology was addressed by grouping proteins that had the same set of peptides to account for their identification. A master protein within a group was assigned based on percent coverage. **Supplementary File: Table S8** provides raw peptide sequences, metrics (retention times, mass, ion scores) and raw abundance values for the 16 total UPLC-MS/MS analyzed samples.

Prior to imputation, a peptide filter was deployed which required a peptide to have a measurable quantitative value in >50% of the samples within a treatment group. After filtering, any missing values were imputed using the following rules: 1) if only a single signal was missing within the group of three, an average of the other two values was used, or 2) if two out of three signals were missing within the group of three, a randomized intensity within the bottom 2% of the detectable signals was used. To summarize the protein abundance, all peptides belonging to the same protein were summed into a single intensity. This consolidation enhances the overall representation of protein abundance, contributing to a more comprehensive analysis. The protein abundance data was then normalized using a robust mean normalization, in which the highest and lowest 10% of the signals were excluded and the average of the remaining signals were normalized to be the same across each individual channel.

In order to assess technical reproducibility, % coefficient of variation (%CV) was calculated for ADH spiked into each sample prior to any type of data normalization. The variation was 19.0%, which is well within expected for non-normalized spiked control data. Then, %CV was calculated for each protein across four injections of the SPQC pool that were interspersed throughout the study. The mean %CV of the SPQC pools was 9.0% which is well within expected analytical tolerances for this type of analysis. To assess biological plus technical variability, %CVs were calculated for each protein, and mean %CV were calculated to be 29.2%, 20.4%, 17.4% and 21.8% for NV-3-L1, V-3-L1, NV-48-L2, and V-48-L2, respectively. Overall, these variations are well within the desired variations and indicate consistent protein extraction, processing and data acquisition.

### 3.5. Identification and annotation of virus-mediated, differentially abundant proteins (vDAPs)

A two-tailed heteroscedastic *t*-test was employed to identify vDAPs in V guts compared to NV guts based on normalized protein abundance and using the criteria of *p*-value < 0.05 and fold change ≥ 2. Amino acid sequences of all vDAPs, i.e., non-redundant vDAPs identified across both larval developmental stages, were subjected to Blastp to query the NCBI non-redundant protein database followed by mapping and annotation with Gene Ontology (GO) terms (E < 1.0E-5) for three provisional functional GO categories (biological process, molecular function, and cellular component) using OmicsBox v2.0 (Götz et al., 2008). To further discriminate more specific putative roles, the vDAP amino acid sequence (one file containing multiple fasta-formatted sequences) were subjected to STRING analysis (https://string-db.org/) (Szklarczyk et al., 2023) to map vDAPs to the program-supplied, annotated *Drosophila melanogaster* proteome in order to predict functional and physical protein networks and to identify significantly enriched functions represented by the vDAPs.

### 3.6. Weighted gene co-expression network analysis (WGCNA)

A matrix of log_2_-transformed, normalized abundance values for all detected proteins (4,189) across the 12 gut samples were subjected to WGCNA for identifying modules (clusters) of co-expressed proteins and their statistical associations with an external sample trait, in this case, TSWV treatment (Langfelder and Horvath, 2008). The adjacencies between proteins were calculated using a soft-thresholding power of 14 under the ‘signed’ network. The adjacency matrix was transformed into a topological overlap matrix for calculating the corresponding dissimilarity. Modules of co-expressed proteins were identified by applying the ‘dynamic tree cut’ method with minimum module size set as 35. By virtue of including all gut proteins identified in the analysis, modules have the potential to include both vDAPs (TSWV-responsive) and non-vDAPs (not responsive to TSWV as compared to NV treatment). Similar modules were merged if their eigengenes shared at least 65% similarity. The module-trait relationships were analyzed by correlating module eigengenes and the infection status (NV or V) of gut samples. Modules that were significantly associated with TSWV treatment were further analyzed for identifying intramodular hub proteins. Proteins that ranked in the top 10% of both module membership (kME, correlation of a protein to its module eigengene) and intramodular connectivity (kIN, sum of connection weights between a protein with all its connected neighbors) among all module proteins were considered as hub proteins. The intramodular interaction networks were visualized with VisANT software v5.53 (Hu et al., 2004). In addition, as described above (section 3.5) for all DAPs, STRING analyses were performed on module sequence sets that were determined to be significantly (*P* < 0.05) associated with the trait of TSWV treatment in order to identify enriched functional categories within modules.

### 3.7. Cross-datasets comparisons of transcriptomes and proteomes of *F. occidentalis* larval guts

Sequence identifiers (OGSv1.1, FOCC IDs) (Rotenberg et al., 2019a, 2019b) of all non-redundant proteins identified in L1 and L2 guts were used to query a *F. occidentalis* larval (L1 and L2) gut transcriptome sequence database assembled previously for a gut RNA-seq experiment (Han and Rotenberg, 2021) that utilized the same insect and virus source, larval developmental stage-time of sample, experimental set-up, conditions and personnel as implemented in the present study. A data matrix consisting of sequence identifiers, sample factors [larval stage (3-L1, 48-L2), treatment (V/NV)], average (n=3) log_2_(normalized transcript abundance) (x) and average (n=3) log_2_(normalized protein abundance) (y) was prepared. Statistical correlation coefficients (Pearson, r) were calculated, and linear regression analyses were performed on all x, y data (all proteins identified with a transcript match) and vDAPs only for each larval stage – treatment combination to determine the strength and significance of the relationship between protein and transcript abundance. Spearman rank correlation coefficients (⍴) were calculated for the sets of x, y data associated for each WGCNA module of co-expressed proteins to examine the possible association between enriched functions ascribed to a module and the strength of the association between protein-transcript abundance. In addition, the list of TSWV-responsive transcripts identified in larval guts previously (105 for 3-L1 and 49 for 48-L2) and gut DAPs identified here were examined for overlap. All correlation and regression analyses were performed with JMP Pro v.15.

## Author Contributions

Conceptualization, D.R and J.H..; methodology, D.R. and J.H.; formal analysis, J.H. and D.R; investigation, J.H.; resources, D.R.; data curation, J.H.; writing-original draft preparation and editing, J.H. and D.R.; visualization, J.H.; supervision, D.R.; project administration, D.R.; funding acquisition, D.R.. All authors have read and agreed to the published version of the manuscript.

## Data Availability Statement

The raw mass spectrometry datasets generated in this study have been deposited in the MassIVE repository (https://massive.ucsd.edu/ProteoSAFe/static/massive.jsp) under project ID: MSV000096811 (available upon publication).

## Acknowledgements

We thank Dr. Erik Soderblum and his staff at the Duke University School of Medicine, Proteomics and Metabolomics Core Facility, for providing consultation and state-of-the-art services of sample processing, protein identification and normalization, and statistical analysis of differential protein expression. We thank Swapna Priya Rajarapu for her assistance with virus acquisition assays and discussions about best practices for thrips dissection for proteomics and data analysis. We also thank Drs. Marlonni Maurastoni and Anna Whitfield for internal review of the manuscript.

