## Supplementary figures and images for "Multi-omics analysis reveals discordant proteome and transcriptome responses in larval guts of *Frankliniella occidentalis* infected with an orthotospovirus"

### Figure S1

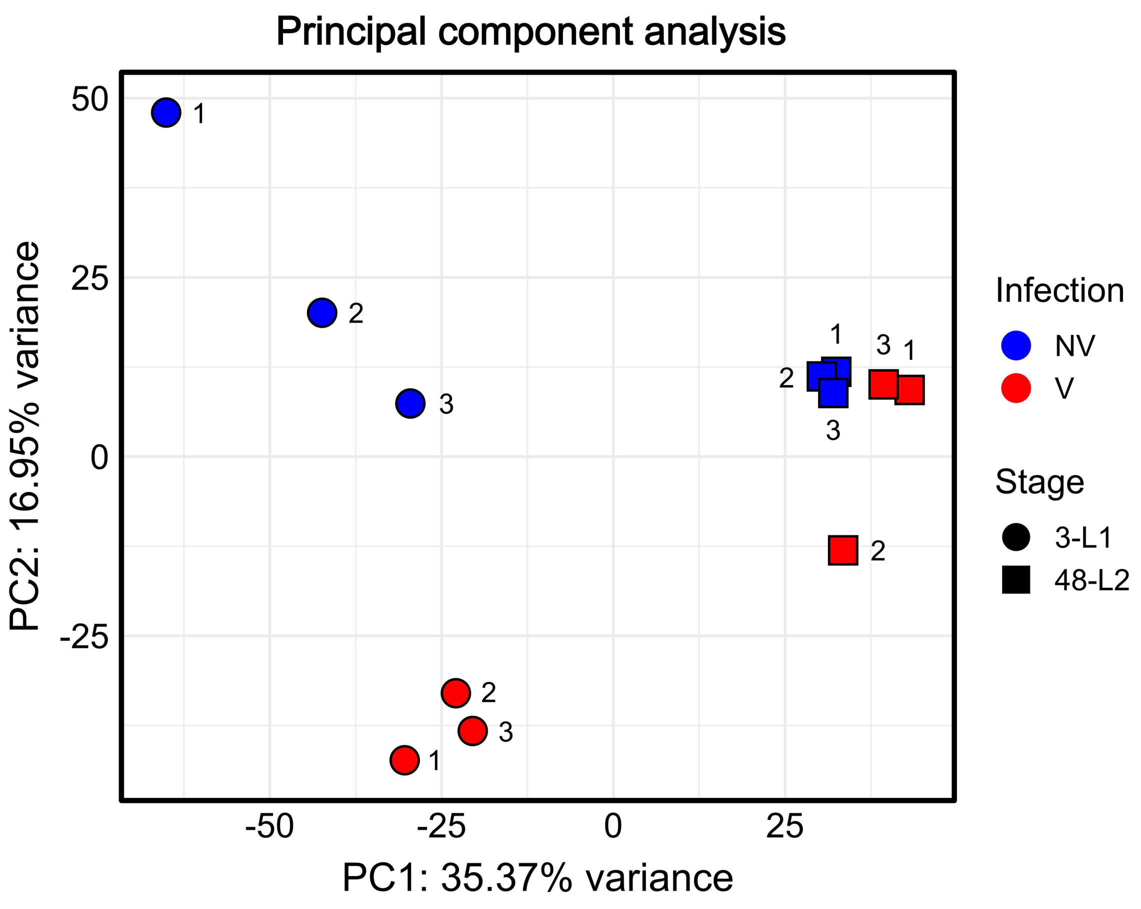

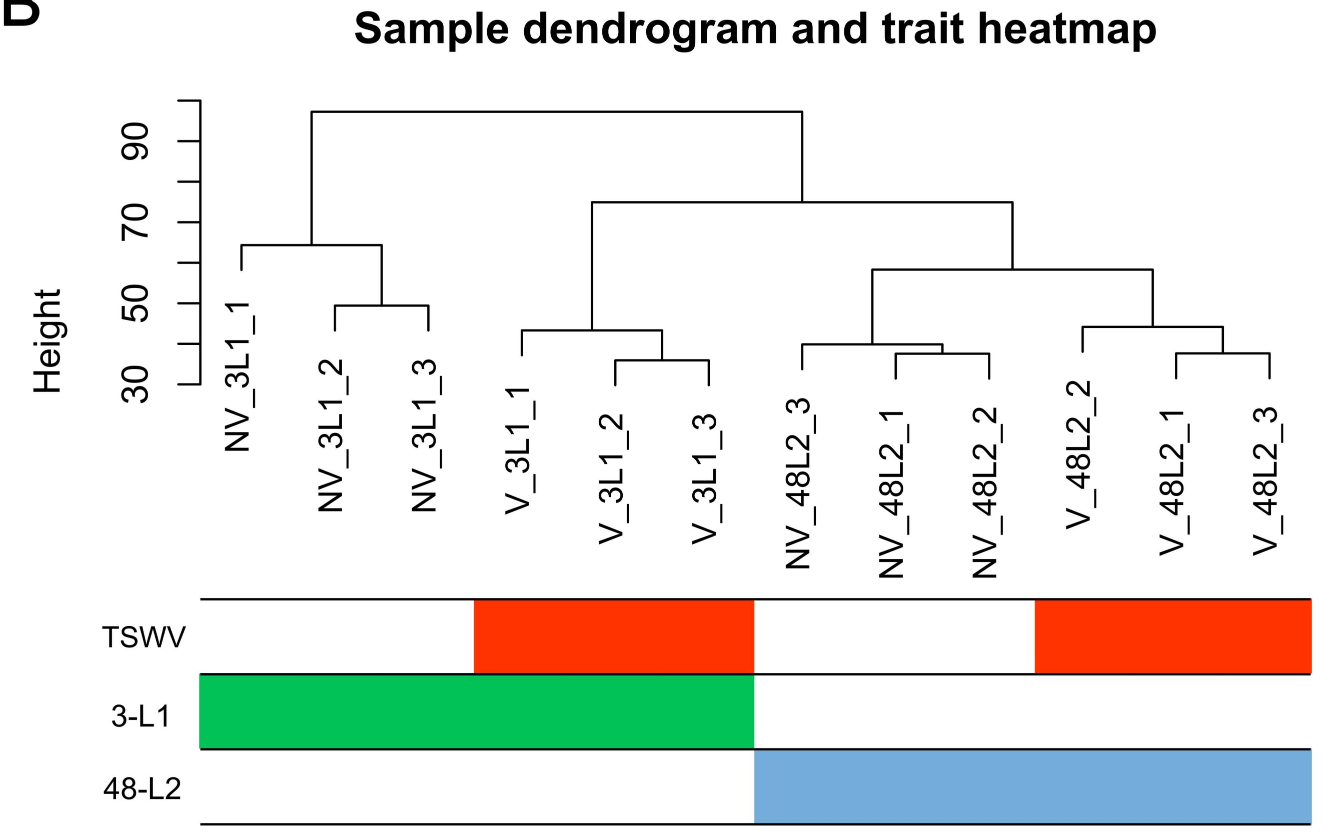

### Figure S2

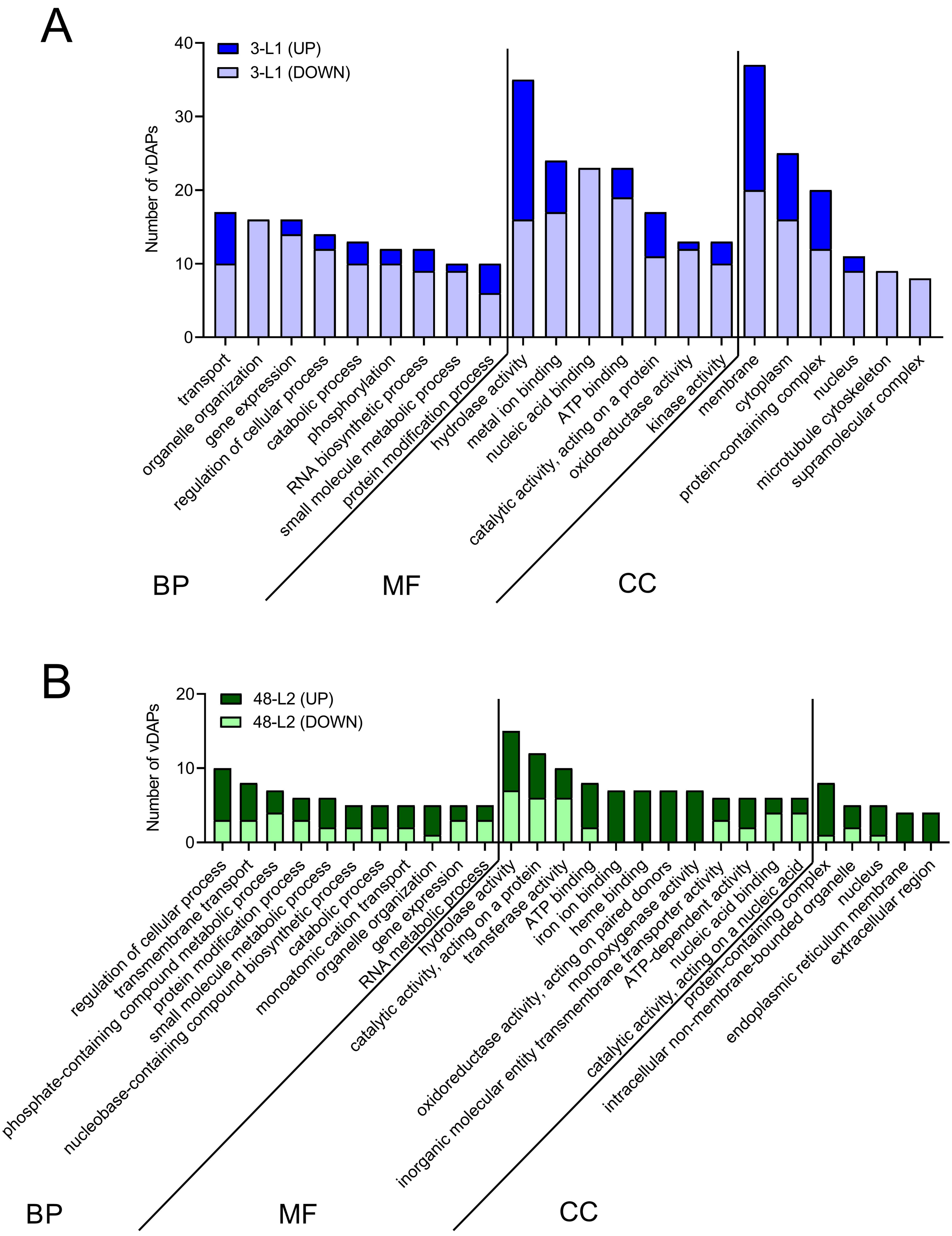

### Figure S3

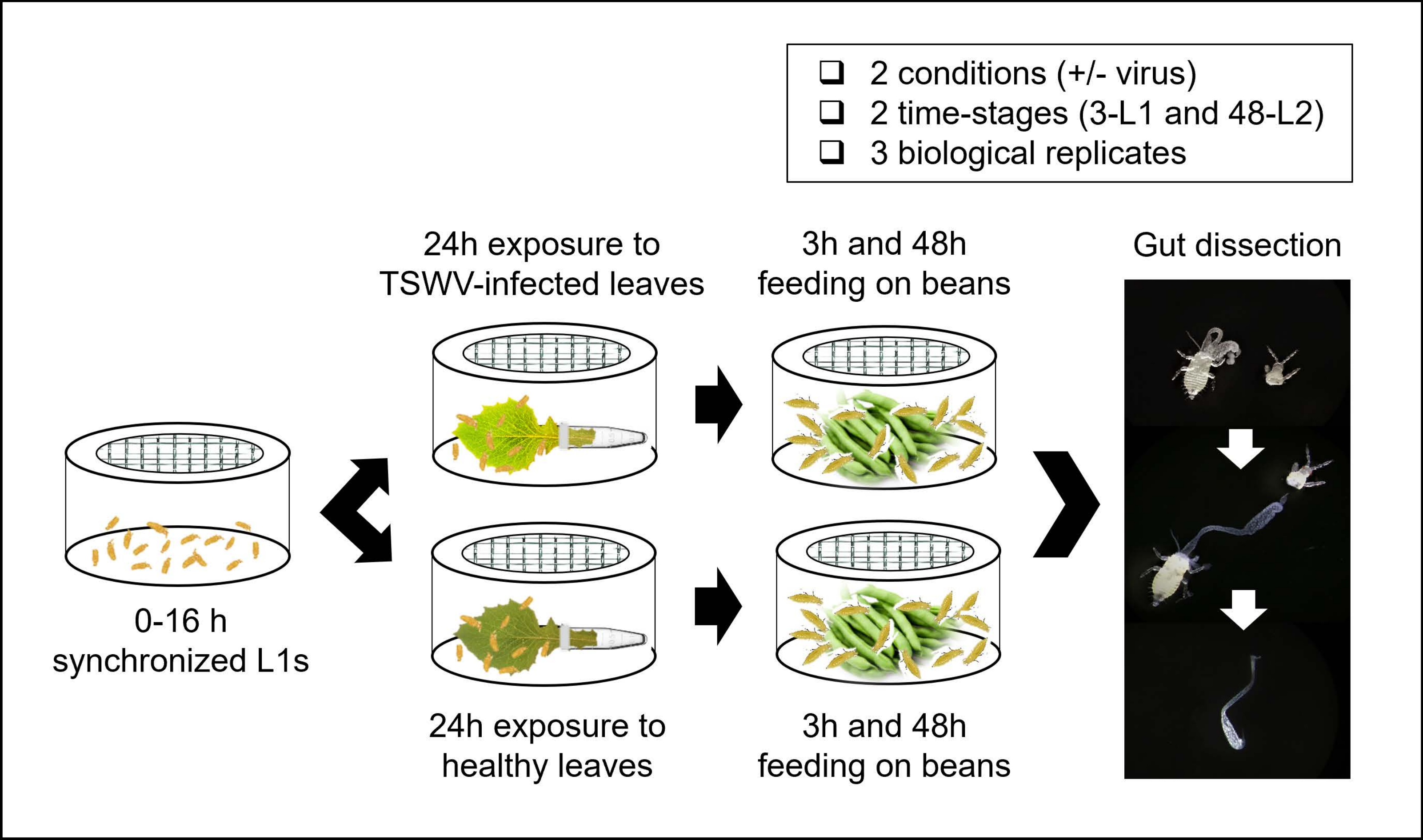
